# NRF2 connects Src tyrosine kinase to ferroptosis resistance in glioblastoma

**DOI:** 10.1101/2023.05.08.539792

**Authors:** Claudia Cirotti, Irene Taddei, Claudia Contadini, Gerardo Pepe, Marco De Bardi, Giovanna Borsellino, Manuela Helmer-Citterich, Daniela Barilà

## Abstract

Glioblastoma (GBM) is a severe brain tumor characterized by an extremely poor survival rate of patients. GBM cancer cells escape to standard therapeutic protocols consisting of combination of ionizing radiation (IR) and alkylating drugs that trigger DNA damage, by rewiring of signaling pathways. In recent years, the upregulation of factors that counteract ferroptosis has been highlighted as a major driver of cancer resistance to IR, although the molecular connection between the activation of oncogenic signaling and the modulation of ferroptosis has not been clarified yet.

Here we provide the first evidence for a molecular connection between the constitutive activation of tyrosine kinases and resistance to ferroptosis. Src tyrosine kinase, a central hub on which Receptor Tyrosine Kinases deregulated signaling converge in cancer, leads to the stabilization and activation of NRF2 pathway, thus promoting resistance to IR-induced ferroptosis. These data suggest that the upregulation of Src-NRF2 axis may represent a vulnerability for combined strategies that, by targeting ferroptosis resistance, enhance radiation sensitivity in glioblastoma.

## INTRODUCTION

Glioblastoma (GBM) is the most aggressive primary brain tumor in adults characterized by poor prognosis (Ostrom *et al*, 2015) linked to radiation-and chemotherapy resistance. Recent studies reported that failure of radiation therapy is also linked to the ability of cancer cells to counteract ferroptosis (Chen *et al*, 2021). Ferroptosis is an iron-dependent type of regulated cell death triggered by disproportionate lipid peroxidation, whose alteration is involved both in tumor development and in response to therapy (Chen *et al*, 2021). The molecular mechanisms that allow cancer cells to overcome ferroptosis have been only partially elucidated.

NRF2 transcription factor, a master regulator of oxidative stress, controls the basal and inducible expression of more than 200 target genes involved in redox homeostasis, metabolism, DNA repair, cell survival and proliferation (Rojo de la Vega *et al*, 2018). NRF2 has a dual role in cancer, being both involved in counteracting cancer initiation and in promoting cancer progression and resistance to therapy (Wu *et al*, 2019). Remarkably, NRF2 upregulation in cancer can successfully face reactive oxygen species (ROS) generated by radio and chemotherapy therefore preventing programmed cell death including ferroptosis (Wang *et al*, 2006; McDonald *et al*, 2010; Ryoo *et al*, 2016; Stockwell *et al*, 2017; Fan *et al*, 2017). NRF2 counteracts ferroptosis by transactivating several cytoprotective genes involved in iron metabolism, ROS detoxification and GSH metabolism (Chen *et al*, 2021). In normal cells, physiological NRF2 turnover, mainly promoted by the KEAP1-CUL3-RBX1 E3-Ubiquitin ligase complex, controls NRF2 protein levels and its activation. Upon stress conditions, ROS modify reactive cysteine residues on KEAP1, preventing its binding to NRF2 which in turn is stabilized (Itoh *et al*, 1999; Tebay *et al*, 2015). Moreover, a non-canonical regulatory pathway involving p62 / sequestosome-1 (SQSTM1) protein has been described (Komatsu *et al*, 2010).

p62 competes with NRF2 for the binding to KEAP1, thus releasing and activating NRF2 (Komatsu *et al*, 2010; Ichimura *et al*, 2013). The molecular mechanisms that allow cancer cells to increase NRF2 expression and therefore dampen ferroptosis, have not been fully elucidated and the interplay between NRF2 and other oncogenes deserves further elucidation. Tyrosine kinases (TKs) signaling emerged as a master class of oncogenes (Blume-Jensen & Hunter, 2001; Yang *et al*, 2022) and represent a core oncogenic requirement for GBM: indeed, up to 50% of GBMs have amplification of a Receptor Tyrosine Kinases (RTKs) followed by the aberrant activation of downstream signaling cascade (Snuderl *et al*, 2011). Src tyrosine kinase is a main node for RTKs signaling being both constantly activated by these signals and responsible for the propagation of downstream events, frequently culminating on transcription factors deregulation (Blume-Jensen & Hunter, 2001;Cirotti *et al*, 2020; Ahluwalia *et al*, 2010).

Here, we uncover NRF2 as a new molecular player downstream Src deregulation in glioblastoma and we highlight an unrevealed link between the aberrant activation of Src and the inhibition of ionizing radiation-induced ferroptosis. We also demonstrate that combining Src targeting with ferroptosis induction significantly increases IR-sensitivity.

## RESULTS

### Constitutive Src kinase activity promotes NRF2 overexpression in glioblastoma

NRF2 transcription factor is aberrantly activated in several types of cancer, sustaining tumor progression, impairing ferroptosis and leading to resistance to chemo-and radiotherapy (Wu *et al*, 2019). A gene expression profile analysis performed by using GEPIA2 (Gene Expression Profile Interactive Analysis 2) web server (Tang *et al*, 2019) highlighted increased NRF2 expression in glioblastoma (GBM), low grade glioma (LGG), pancreatic adenocarcinoma (PAAD) and thymoma (THYM) (**Figure 1A**). Focusing on GBM we next compared the expression of NRF2 gene between tumor samples from TCGA and normal brain tissues, and we observed a significant increased expression of NRF2 in tumors (**Figure 1B**). We next moved to *in vitro* GBM cellular models taking advantage of U87-MG and T98G cell lines and of GBMSC83 neurospheres derived from primary GBM tumors (Minata *et al*, 2019; Mao *et al*, 2013; Contadini *et al*, 2023).

**Figure 1.**
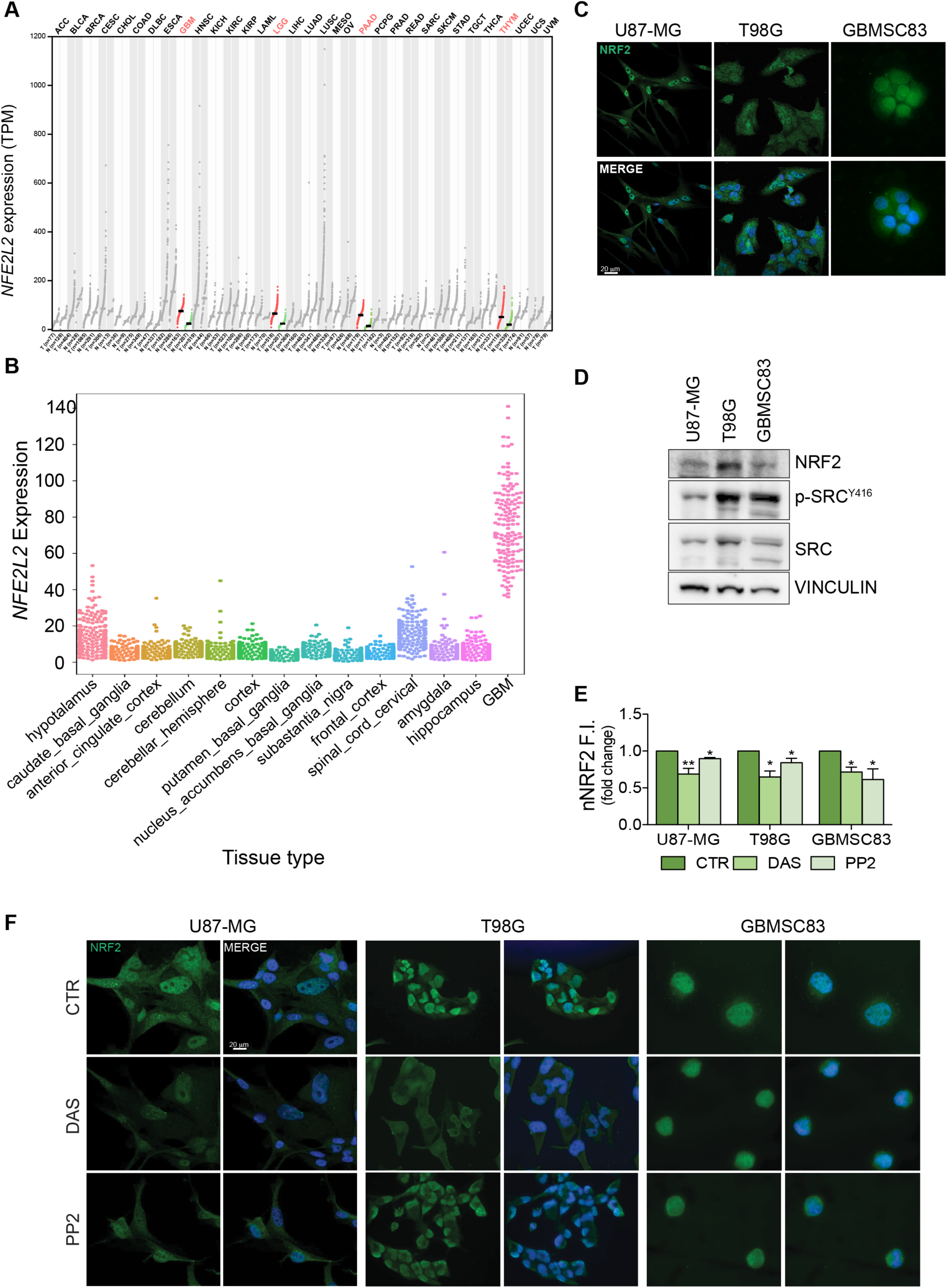
NRF2 is highly expressed in glioblastoma and it is sustained by Src tyrosine kinase activity. **A**) *NFE2L2* Gene Expression Profile across tumor samples and paired normal tissues (Dot plot; each dot represents *NFE2L2* expression in each sample (ANOVA; match TGCA normal and GTEx data); **B**) *NFE2L2* gene expression in glioblastoma (TCGA-GBM) compared to normal brain areas (GTEx); **C**) Immunofluorescence of U87-MG, T98G and GBMSC83 cells; NRF2 (green); DNA (Hoechst, blue). **D**) Immunoblotting of NRF2, p-Src^Y416^ and Src in U87-MG, T98G and GBMSC83 cells. Vinculin was used as loading control. **E-F**) Immunofluorescence experiments of U87-MG, T98G and GBMSC83 cells upon 16hrs treatments with Dasatinib (DAS) 10nM (U87-MG, T98G) and 1μM (GBMSC83); PP2 5μM. **E**) Histogram of NRF2 nuclear (nNRF2) fluorescence intensity (F.I.); Statistical analyses: paired Student’s t-test: (* p < 0.05; ** p < 0.01). **F**) Representative immunofluorescence of U87-MG, T98G and GBMSC83 cells treated as before. NRF2 (green); DNA (Hoechst, blue).

Immunofluorescence analyses confirmed that NRF2 is abundantly expressed and localized in the nuclear compartment in all the GBM cellular models (**Figure 1C**). Tyrosine kinases deregulation is among the most studied pro-tumoral factor in glioblastoma and Src tyrosine kinase in this tumor is constantly activated (Bjorge *et al*, 2000; Du *et al*, 2009). Immunoblotting with anti-pSrc^Y416^ antibodies confirmed that the constitutive activation of Src in our GBM models (**Figure 1D**). Given the simultaneous aberrant activation of Src kinase and the increased levels of NRF2, we tested whether the pharmacological targeting of Src kinase activity with Dasatinib and PP2 may affect NRF2 expression.

Interestingly, immunofluorescence experiments show that the inhibition of Src (**Supplementary Figure S1)** significantly decreased NRF2 nuclear localization (**Figures 1E-F**). Furthermore, NRF2 localization was evaluated also upon subcellular fractionation. Pharmacological Src kinase inhibition (**Supplementary Figure S2**) significantly impinged on NRF2 nuclear localization, similarly to what observed when cells were treated with Trigonelline, a well-known NRF2 inhibitor (Arlt *et al*, 2013) (**Figure 2A)**. Confocal microscopy analyses further confirmed that Dasatinib significantly affected NRF2 nuclear localization, resulting in a reduced ratio of nuclear/cytoplasmic NRF2 fluorescence intensity (**Figure 2B**). To unambiguously demonstrate that Src may modulate NRF2, Src kinase activity was also genetically modulated taking advantage of Src mutants that either mimic the constitutive activation (Src^Y527F^) or the enzymatic inactivation (Src^K295M^). Interestingly, the overexpression of Src^K295M^ caused a decrease of NRF2 protein expression (**Figure 2C)** and nuclear localization (**Figure 2D**). We next generated GBM cell lines stably overexpressing *wild-type* Src (Src^WT^) or the catalytically inactive mutant Src^K295M^ (hereafter named Src Kinase-Dead, Src^KD^) in which we could once again recapitulate that Src activity is required to sustain NRF2 (**Figure 2E-F**). Overall, these data allow the conclusion that Src activity sustains NRF2 expression and nuclear localization in GBM cells.

**Figure 2.**
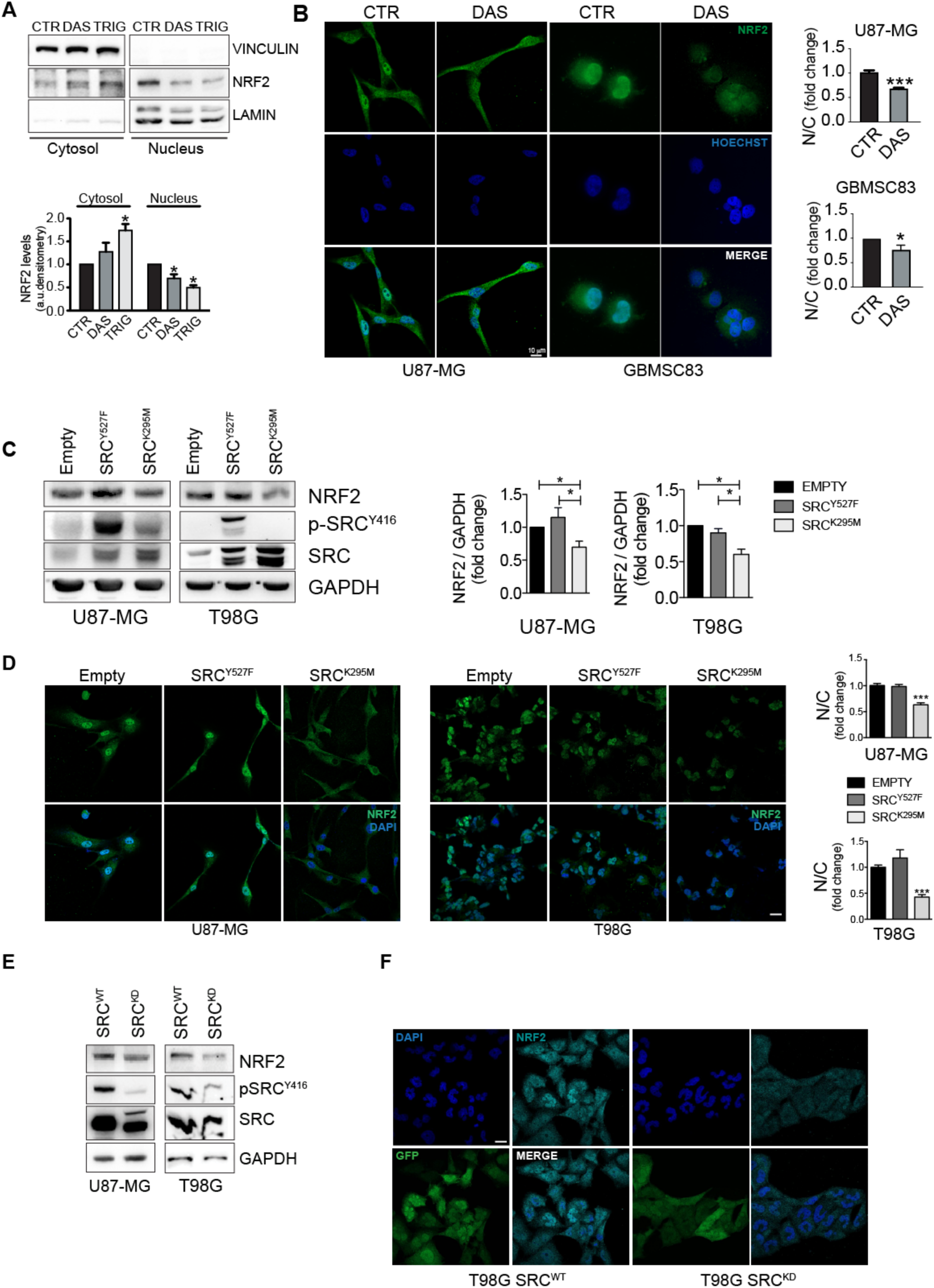
Pharmacological and genetic modulation of Src activity affects NRF2 expression. **A**) Immunoblotting and relative densitometric analysis of NRF2 in U87-MG cytosolic and nuclear cellular fractions, after DAS 10 nM or Trigonelline (Trig) 5 μM treatment for 6hrs. **B**) Confocal Immunofluorescence experiments and relative quantifications in U87-MG and GBMSC83 cells treated for 16hrs with DAS 10 nM and 1μM, respectively. NRF2 (green) and DNA (Hoechts, blue); **C**) Immunoblotting and relative densitometric analysis of NRF2, p-Src^Y416^ and Src, and **D**) Immunofluorescence and relative quantification in U87-MG and T98G cells after 24hrs of transient tranfections with active Src (Src^Y527F^) or catalytically inactive mutant (Src^K295M^). NRF2 (green); DNA (Hoechst, blue). **E**) Immunoblotting of NRF2, p-Src^Y416^ and Src in U87-MG and T98G stably overexpressing *wild-type* Src (Src^WT^) or the catalytically inactive mutant (Src^K295M^, hereafter Src^KD^). **F**) Immunofluorescence analysis of T98G Src^WT^ and Src^KD^ cells. NRF2 (Cyan); GFP (green); DNA (Hoechts, blue). Statistical analyses: paired Student’s t-test (* p < 0.05; *** p < 0.001).

### Src kinase activity promotes NRF2 signaling

To assess whether Src kinase may modulate NRF2 functionality, NRF2 transcriptional activity was evaluated upon pharmacological or genetic modulation of Src. Remarkably, Dasatinib treatment triggered the downregulation of several NRF2 target genes, namely *heme oxygenase 1* (*hmox1*), *glutamate-cysteine ligase catalytic subunits* (*c-gcl*), *NAD(P)H Quinone Dehydrogenase 1* (*nqo1*) and *sequestosome-1* (*sqstm1*), as revealed by qRT-PCR, both in U87-MG cells and in GBMSC83 neurospheres (**Figure 3A**). Consistently immunoblotting analyses revealed a significant decrement of SQSTM1/p62 (hereafter, p62) and heme oxygenase 1 (HO-1) proteins upon Dasatinib treatment (**Figure 3B**). The transcriptional activity of NRF2 was also significantly decreased in U87-MG and T98G cells engineered to stably express Src^KD^, compared to the ones expressing Src^WT^ counterpart (**Figure 3C** and **3D**) as well as upon transient transfection of the catalytically inactive in Src^K295M^ (**Supplementary Figure S3)**. Overall, these data suggest that Src activity sustains NRF2 transcriptional activity.

**Figure 3.**
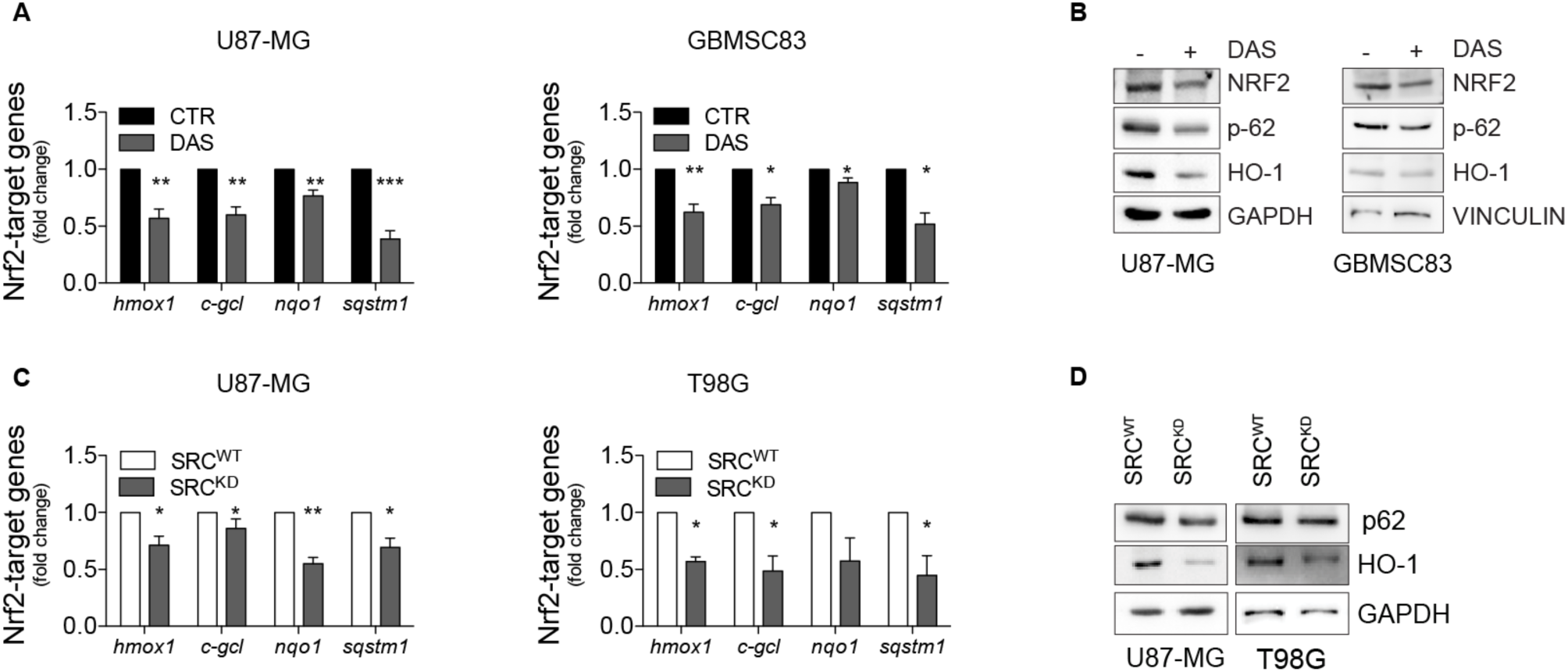
Pharmacological and genetic modulation of Src activity affects NRF2 transcriptional activity. **A**) RT-qPCR of NRF2 target genes (Actin: housekeeping gene); and **B**) Immunoblotting of NRF2, p62 and HO-1 in U87-MG and GBMSC83 cells treated for 16hrs with DAS 10nM and 1μM, respectively. **C**) RT-qPCR of NRF2 target genes (Actin: housekeeping gene); and **D**) Immunoblotting of p62 and HO-1 in U87-MG and T98G cells stably overexpressing Src^WT^ or Src^KD^. GAPDH was used as loading control. Statistical analyses: paired Student’s t-test (* p < 0.05; ** p < 0.01 ; *** p < 0.001).

### Src kinase activity inhibition turns-off NRF2 pathway affecting p62-KEAP1 interaction

Previous work highlighted a strong correlation between NRF2 expression and p62 in GBM tumors, and demonstrated that the high expression levels of NRF2 protein may be sustained by the ability of p62 to bind KEAP1 preventing its interaction with NRF2 (Pölönen *et al*, 2019). We therefore asked the question whether the aberrant activity of Src may sustain p62-KEAP1 interaction therefore enhancing NRF2 expression and activity. Confocal microscopy analysis on GBM cells treated or not with Dasatinib, highlighted the presence of p62 aggregates and a strong co-localization between p62 and KEAP1 in these structures (**Figure 4A-B**). More interestingly, Dasatinib treatment significantly released p62-KEAP1 interaction and promoted a diffused relocalization of KEAP1 into the cytosolic compartment, particularly in the perinuclear area (**Figure 4A-B**). The competitive interaction between p62 and KEAP1 is ensured by the phosphorylation of p62 on Serine 349 residue (S349) (Ichimura *et al*, 2013). Interestingly, Dasatinib treatment, severely impinged on this phosphorylation (**Figure 4C**), suggesting that Src activity could indirectly promote p62 phosphorylation on S349 which drives the formation of KEAP1-p62 complex, ultimately leading to NRF2 stabilization and accumulation. Confocal microscopy analyses on Src^WT^ or Src^KD^ cells further confirmed this hypothesis (**Figure 4D-E**). Furthermore, Src^KD^ triggered the reduction of p62 phosphorylation on S349 (**Figure 4F**), similarly to what observed with Dasatinib. These data demonstrated that Src kinase activity sustains p62-KEAP1 interaction therefore regulating NRF2 functionality.

**Figure 4.**
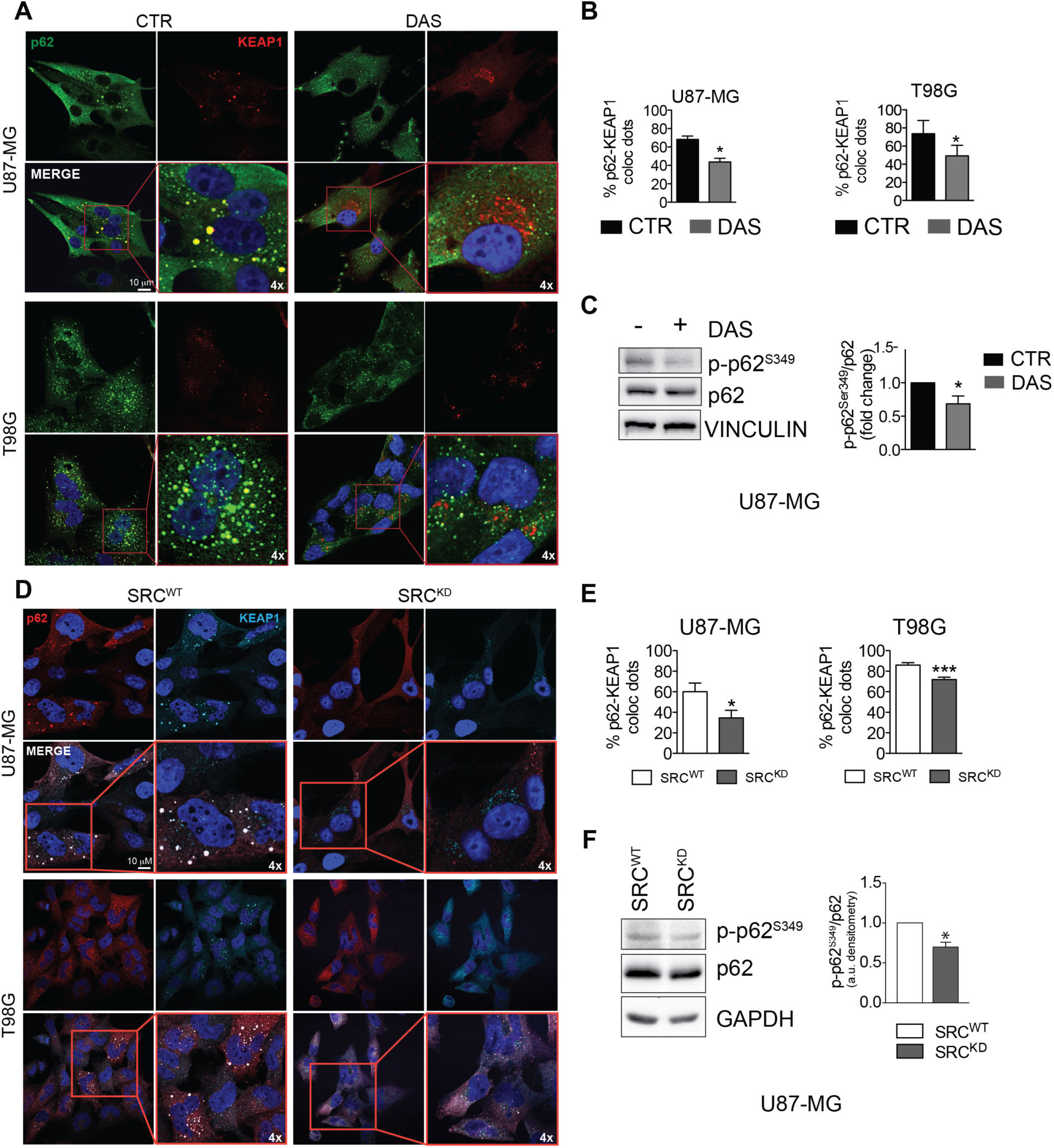
Src activity sustains p62-KEAP1 interaction promoting p62^S349^ phosphorylation. **A-B**) Confocal microscopy analyses (**A**) and quantification of colocalizing dots (**B**) in U87-MG and T98G cells treated with DAS 10nM for 16hrs. p62 (green); KEAP1 (red); DNA (Hoechst, blue); 4x digital magnification showing merged signals. **C**) Immunoblotting and relative densitometric analysis of p-p62^S349^ and p62 in U87-MG cells treated as previously. **D-E**) Confocal microscopy analyses (**D**) and quantification of colocalizing dots (**E**) in U87-MG and T98G cells stably overexpressing Src^WT^ or Src^KD^. p62 (red); KEAP1 (cyan); DNA (DAPI, blue); 4x digital magnification showing merged signals. **F**) Immunoblotting and relative densitometric analysis of p-p62^S349^ and p62 in U87-MG cells stably overexpressing Src^WT^ or Src^KD^. Statistical analyses: paired (**C**, **F**) or unpaired (**B**, **E**) Student’s t-test: (* p < 0.05; ** p < 0.01; *** p< 0.001).

### Src kinase activity inhibition promotes NRF2-KEAP1 interaction triggering NRF2 degradation

The observation that the inhibition of Src activity leads to the release of KEAP1-p62 interaction supported the hypothesis that in this context Src inhibition may conversely promote the interaction between KEAP1 and NRF2. Importantly, we could show that Src^K295M^ transient transfection drove the co-localization between KEAP1 and NRF2 (**Figure 5A,** red boxes -4x magnification). Of note, we immunoprecipitated KEAP1 from U87-MG Src^WT^ and Src^KD^ cells and we observed that the interaction between KEAP1 and NRF2 is increased in Src^KD^ cells, despite of the decreased levels of NRF2 protein in this condition (**Figure 5B**). Consistently, Src^KD^ overexpression decreased NRF2 protein stability **(Figure 5C)**. Altogether these data strongly support the conclusion that Src kinase activity enhances NRF2 protein stability.

**Figure 5.**
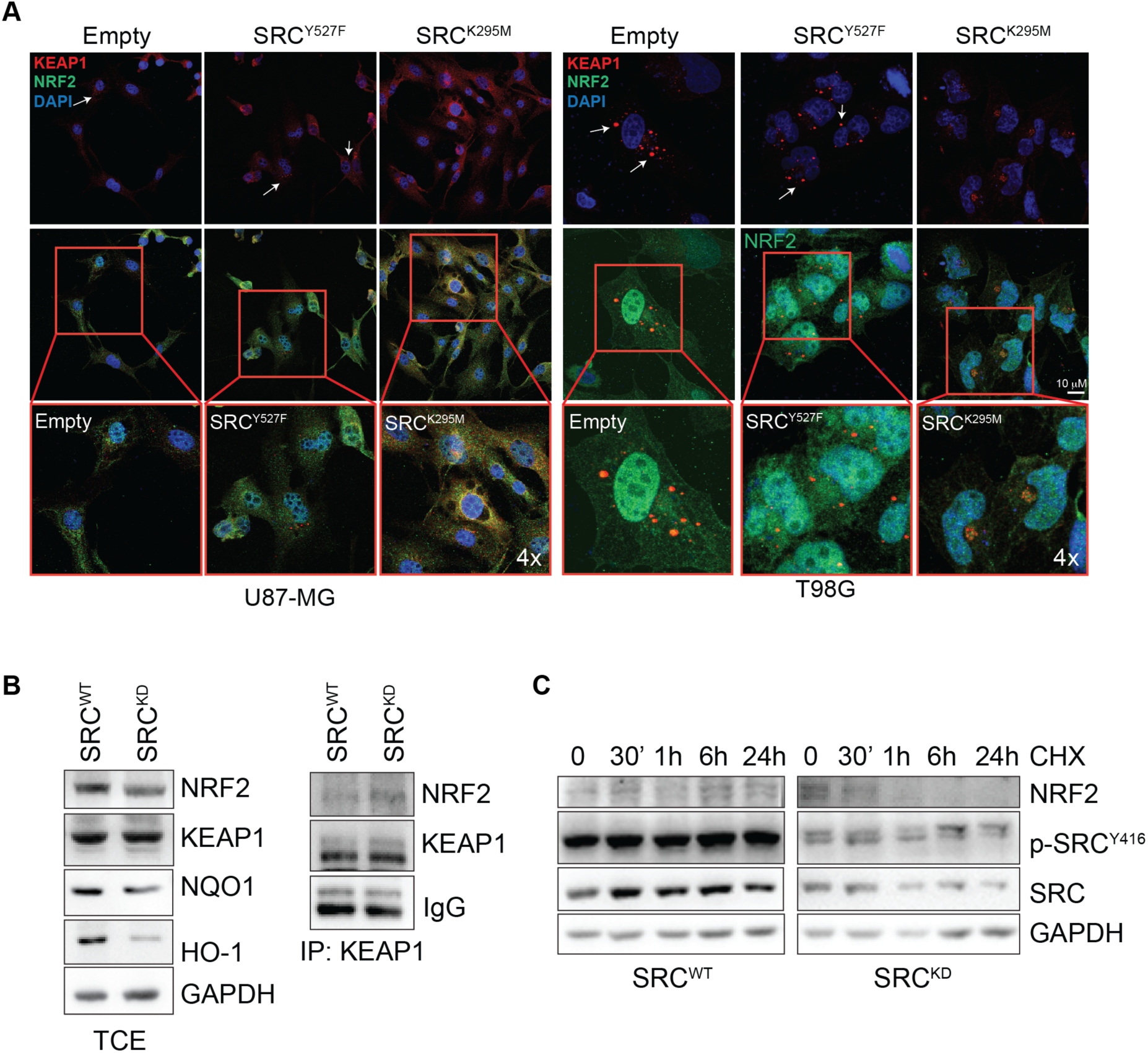
Src activity affects KEAP1 intracellular distribution and its interaction with NRF2. **A**) Confocal microscopy analyses in U87-MG and T98G cells after 24hrs of transient tranfections with active Src (Src^Y527F^) or catalytically inactive mutant (Src^K295M^). NRF2 (green); KEAP1 (red); DNA (Hoechst, blue). 4x digital magnification showing merged signals. **B**) Co-immunoprecipitation experiments of KEAP1 in U87-MG cells stably overexpressing Src^WT^ or Src^KD^ revealing KEAP1-NRF2 interaction. **C**) Immunoblotting of NRF2, p-pSrc^Y416^ and Src in U87-MG Src^WT^ and Src^KD^ treated with Cycloheximide (CHX) 100μg/ml for 0, 30 min, 1, 6 and 24hrs. GAPDH was used as loading control.

### Src activity sustains mTORC1-dependent p62 phosphorylation

It has been previously reported that the competitive binding between p62 and KEAP1 proteins is ensured by mTORC1-dependent phosphorylation of p62 on serine 349 residue (Ichimura *et al*, 2013). Data from literature also support a direct role for Src kinase in modulating mTORC1 activity in several type of cancers (Pal *et al*, 2018; Vojtěchová *et al*, 2008), raising the question whether the constitutive activity of Src may trigger the aberrant activation of mTORC1 in glioblastoma. Immunoblotting experiments revealed the presence of phosphorylated p70S6K (p-p70S6K^T389^), a well-known target of mTORC1 (Maiese *et al*, 2013) in our cellular models (**Figure 6A**). Remarkably, pharmacological and genetic targeting of Src kinase activity resulted in the downregulation of p70S6K phosphorylation(**Figure 6A**). To further confirm the involvement of mTORC1 in sustaining Src-dependent NRF2 hyperactivation, we took advantage of the mTORC1 inhibitor Rapamycin. As expected, Rapamycin turned off mTORC1 activity, as shown by the downregulation of p-p70S6K^T389^ (**Supplementary Figure S4**). Of note, NRF2-dependent pathway was strongly affected by Rapamycin, as shown by the decreased expression levels of NRF2 and of its target genes. In addition, Rapamycin treatment dampened p62 phosphorylation on S349, thus impairing KEAP1 sequestration (**Figure 6B** and **Supplementary Figure S5-S6**). Overall, these data suggest that Src may depend on mTORC1 activation to sustain p62 phosphorylation and NRF2 hyperactivation in glioblastoma cellular models. Interestingly, comparing the expression profiles of glioblastoma patients (TGCA data) with normal brain tissues (GTEx data) we found a significant up-regulation in tumor samples of several genes enriched mTORC1 pathway, thus strengthening the relevance of this hallmark in glioblastoma (**Supplementary Figure S7**).

**Figure 6.**
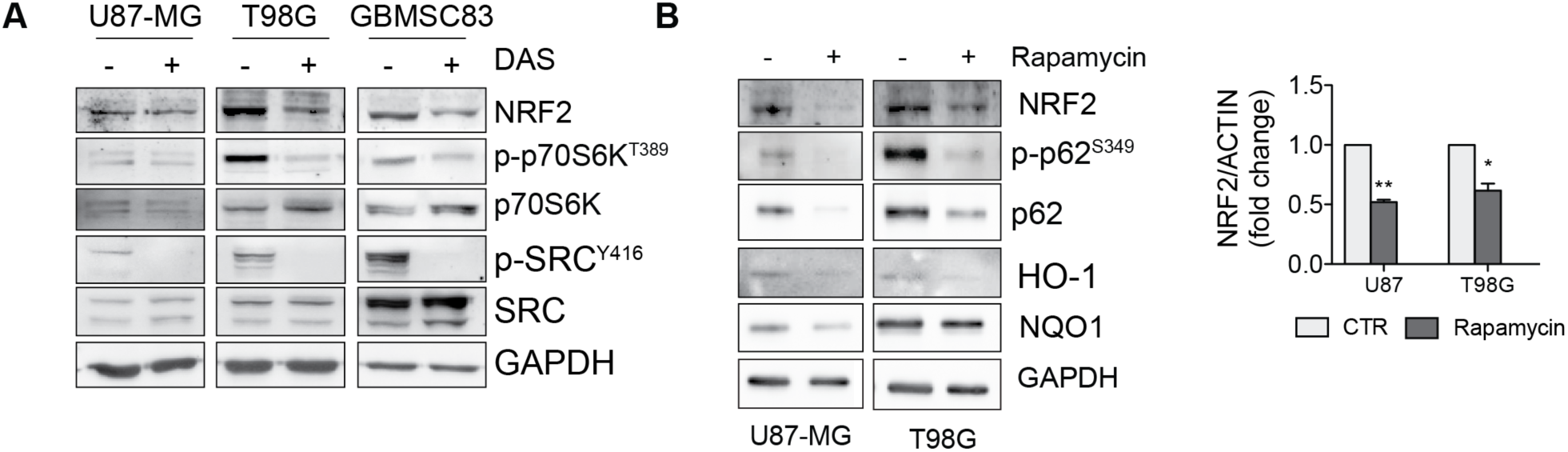
Src activity sustains mTORC1-dependent signaling. **A**) Immunoblotting of p-p70S6K^T389^ and p70S6K in U87-MG, T98G and GBMSC83 cells treated for 16 hrs with DAS 10nM and 1μM, respectively. **B**) Immunoblotting of NRF2, p-p62^S349^, p62, HO-1 and NQO1 in U87-MG and T98G cells upon Rapamycin 100nM treatment for 48hrs, and relative NRF2 densitometric analysis.

### NRF2 is required for Src-dependent proliferation, clonogenicity and resistance to ionizing radiation treatment

It has been shown that increased NRF2 protein levels represent a major trigger for cancer resistance to therapy (Awuah *et al*, 2022) NRF2 indeed is strongly involved in tumor growth (Rojo de la Vega *et al*, 2018), it can sustain and promote malignant transformation of GBM stem cells (Zhu *et al*, 2013) and it has been shown to be responsible for chemo-and radiotherapy resistance in glioblastoma cellular models (Rocha *et al*, 2016; Singer *et al*, 2015) shifting resistant cells towards mesenchymal phenotype (Singer *et al*, 2015). We therefore asked whether turning off NRF2 functionality in tumors that upregulate Src activity may enhance the therapeutic response. We could show that NRF2 inhibition by Trigonelline treatment significantly affected Src^WT^, but not Src^KD^, cell proliferation (**Figure 7A**). Clonogenic assay experiments demonstrated that NRF2 inhibition *via* ML385 has a stronger effect on the clonogenic potential of Src^WT^ cells compared to Src^KD^ ones (**Figure 7B**).

**Figure 7.**
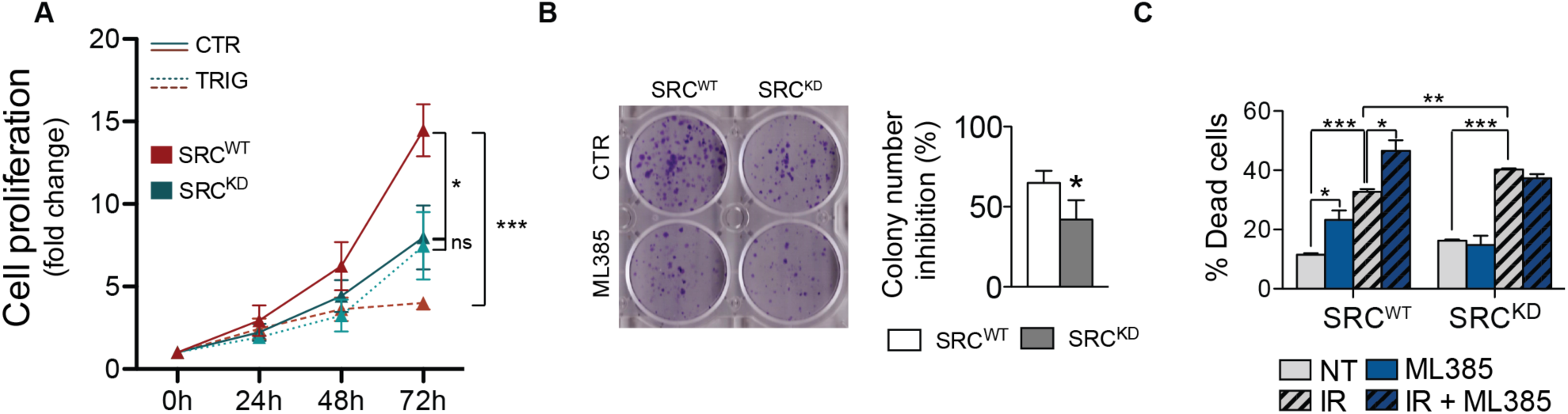
NRF2 inhibition affects cell proliferation and sensitivity to ionizing radiation in SRC^WT^ cells. **A**) Cell proliferation assay in T98G cells stably overexpressing Src^WT^ or Src^KD^ treated or not with Trigonelline 5μM for 0, 24, 48 and 72hrs. **B**) Clonogenic assay performed on T98G cells stably overexpressing Src^WT^ or Src^KD^ treated or not with ML385 5μM and relative histogram representing colony number inhibition (as pecentage to each control condition). **C**) Histogram of the percentage of cell death from cytofluorimetric analysis in T98G cells stably overexpressing Src^WT^ or Src^KD^ 48hrs from irradiation (10Gy), upon pretreatment or not with ML385 5μM. Statistical analyses: unpaired Student’s t-test: (* p < 0.05; ** p < 0.01; *** p< 0.001).

To further strengthen NRF2 role downstream Src activity we evaluated GBM cells sensitivity to ionizing radiation (IR), that represents the standard therapeutic approach for GBM patients. As expected, Src^KD^ cells were slightly but significantly more sensitive to IR than Src^WT^ cells (**Figure 7C**). Of note, NRF2 inhibition significantly affected cell viability in Src^WT^ and not in Src^KD^ cells and significantly increased sensitivity to IR-induced cell death only in Src^WT^ cells (**Figure 7C**).

### Src kinase activity drives NRF2-dependent resistance to ionizing radiation-induced ferroptosis

Recently, a novel role of NRF2 as inhibitor of ferroptosis, has been reported also in GBM (Fan *et al*, 2017; Dodson *et al*, 2019). Importantly, SLC7A11 and GPX4, two genes that play a key role in the modulation of ferroptosis, have been also shown to be NRF2 target genes (Wu *et al*, 2019). We therefore asked the question whether Src-NRF2 axis may also sustain the upregulation of these genes, preventing ferroptosis. Genetic (Src^KD^ cells) and pharmacological (Dasatinib) inhibition of Src kinase activity significantly decreased the expression of SLC7A11 and GPX4 genes (**Figures 8A-B**). Given this data, we sought to investigate the significance of ferroptosis in our experimental models. After 48h from irradiation, we observed a significant reduction in the percentage of cell death when ferroptosis was inhibited by Ferrostatin-1 (Fer-1) (**Figure 8C**), supporting the conclusion that IR-induced cell death occurs also *via* ferroptosis in our cells. The role of tyrosine kinases in the modulation of ferroptosis has not been clearly investigated yet. We asked whether the increased sensitivity to IR detected in Src^KD^ cells (**Figure 7C**) could be due to ferroptosis. Indeed, the inhibition of ferroptosis has a more dramatic effect on Src^KD^ cells compared to the control ones (**Figures 8D**). To strength the significance of our finding we also performed experiments on patient derived GBMSC83 neurospheres. We observed that dasatinib treatment sensitized GBMSC83 to IR. Importantly, cotreatment with Dasatinib and Fer-1 failed to enhance the percentage of cell death (**Figure 8E**), thus suggesting that indeed Src inhibition sensitize cells to ferroptosis. In line with this, lipid peroxidation increased upon Dasatinib and irradiation cotreatment (**Figure 8F**). Importantly, ferroptosis induction by Erastin was strongly induced by the combined treatment with Dasatinib and, more interestingly, this combination strongly sensitized cells to IR (**Figure 8G** and **Supplementary Figure S8**). Altogether these data demonstrate that Src-NRF2 axis sustain cancer cells resistance to ionizing radiation by preventing ferroptosis (**Figure 8H**).

**Figure 8.**
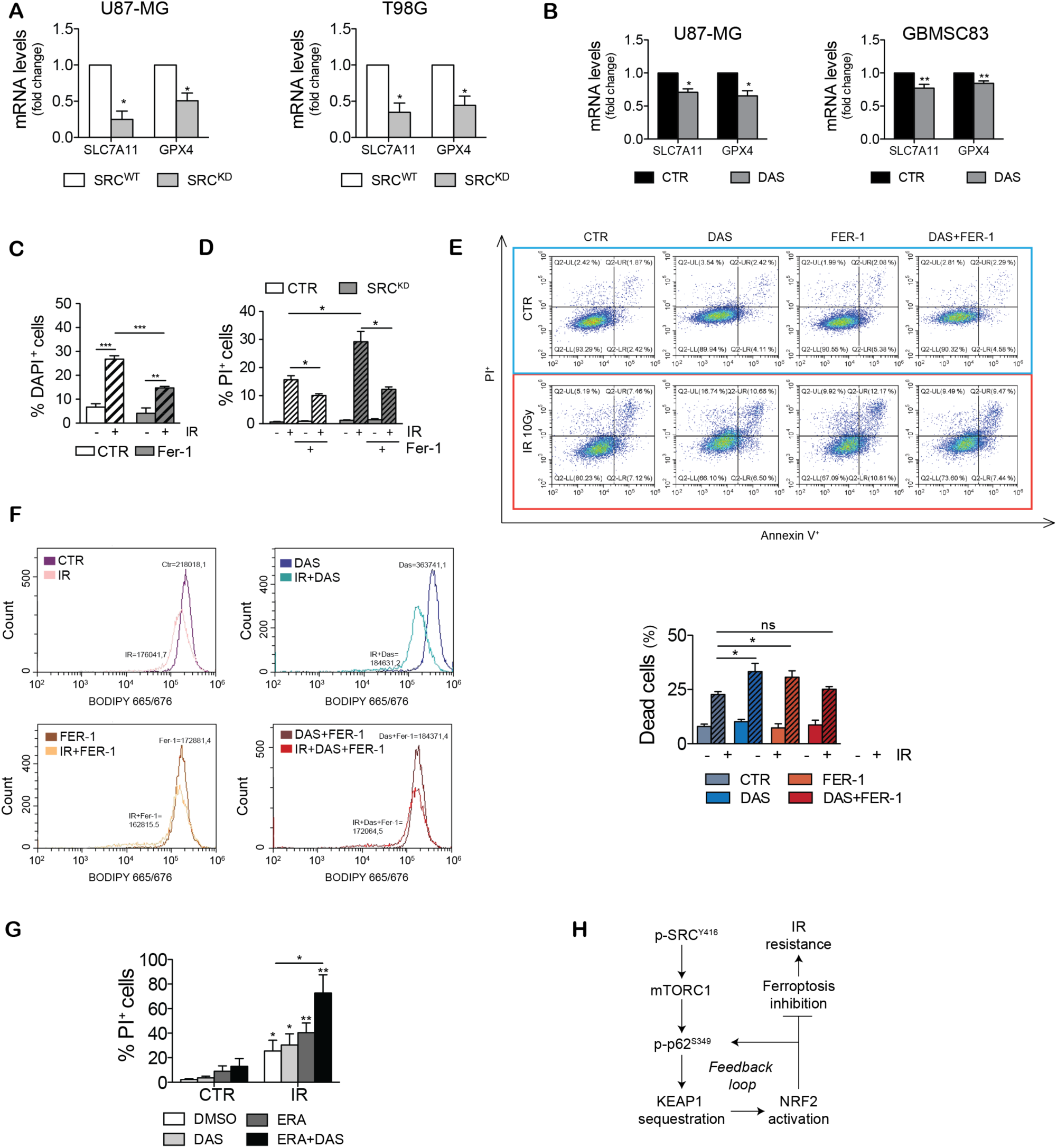
NRF2 inhibition sensitizes GBM cells to ionizing radiation-induced ferroptotic cell death. **A-B**) RT-qPCR of ferroptotic genes *slc7a11* and *gpx4* in U87-MG and T98G stably expressing SRC^WT^ or SRC^KD^ (**A**) and in U87-MG and GBMSC83 cells treated with DAS 10nM and 1μM, respectively (**B**) (Actin: housekeeping gene). **C**) Histogram of DAPI^+^ T98G cells after 48hrs from irradiation (10Gy), with or without Ferrostatin-1 (Fer-1) 5μM pretreatment. **D**) Histogram of Propidium Iodide (PI^+^) T98G SRC^WT^ and SRC^KD^ cells after 48hrs from irradiation (10Gy), with or without Fer-1 5μM treatment. **E**) Cytofluorimetric analysis and relative histogram of cell death upon AnnexinV-PI staining in GBMSC83 cells after 48hrs irradiation (10Gy), with or without DAS 1μM and/or Fer-1 5μM. **F**) Cytofluorimetric analysis of lipid peroxidation upon BODIPY 665/676 staining in in GBMSC83 cells after 48hrs irradiation (10Gy), with or without DAS 1μM and/or Fer-1 5μM. **G**) Histogram of Propidium Iodide (PI^+^) T98G cells after 48hrs irradiation (10Gy), with or without DAS 10nM and/or Erastin (ERA) 1μM. Statistical analyses: paired (**A**,**B**) and unpaired (**C,D,E,G**) Student’s t-test: (* p < 0.05; ** p < 0.01; *** p< 0.001). (**H**) Working model.

## DISCUSSION

Radiation and chemotherapy resistance is a major issue in cancer therapy as cancer cells activate several signaling pathways to counteract cell death induction. Ferroptosis inhibition is emerging as one of the major mechanism responsible for IR resistance (Chen *et al*, 2021) although the molecular mechanisms that allow cancer cells to prevent ferroptosis have been only partially elucidated.

NRF2 transcription factor, a major controller of oxidative stress, has been shown to be aberrantly up regulated in many tumors, including glioblastoma (Rojo de la Vega *et al*, 2018). More importantly, its overexpression can drive cancer resistance to chemo and radiotherapy induced apoptosis and ferroptosis, therefore representing one of the major triggers of poor prognosis in these tumors (Wu *et al*, 2019). Its aberrant activation relies often on NRF2 or KEAP1 gene mutation, both resulting in increased NRF2 protein levels (Kansanen *et al*, 2013). Nevertheless, the upregulation of NRF2 has been also detected independently of these events pointing to alternative signaling molecules iperactivated in cancer as possible NRF2 regulators. Recently, the bioinformatic analysis of gene expression data from the Broad-Novartis Cancer Cell Line Encyclopedia (CCLE) and the Cancer Genome Atlas (TCGA) identified glioblastoma as one of the tumors where NRF2 is more often upregulated (Pölönen *et al*, 2019). We confirmed this issue (**Figure 1**) and we therefore selected GBM cellular models as experimental system to test the hypothesis that other signaling pathways often deregulated in GBM may contribute to increase NRF2 expression levels. Genomic and transcriptomic analyses uncovered RTKs signaling as one of the pathways that are more frequently deregulated in GBM (Cancer Genome Atlas Research Network, 2008). More than 60% of GBM display aberrant activation of EGFR or MET or HER2. Importantly, a common trait of the signaling cascade downstream RTKs is the constitutive activation of Src kinase (REF Du et al., 2009). Based on these evidences we decided to test the hypothesis that Src activity may impact on NRF2 expression and signaling. Here by using GBM cell lines we identified for the first time a role for Src kinase activity as modulator of NRF2. Importantly, we confirmed that in our experimental systems both NRF2 protein expression and Src activity were up-regulated (**Figure 1**) and that both pharmacological and genetic inhibition of Src kinase activity significantly downregulated NRF2 protein expression levels (**Figure 1** and **Figure 2**). Immunofluorescence and subcellular fractionation experiments showed that Src sustains NRF2 localization in the nucleus (**Figure 2**), thus promoting the expression of selected NRF2-target genes related to the oxidative stress response (**Figure 3**) and to ferroptosis (**Figure 8**). We also reported that Src caused the formation of p62 aggregates that colocalize with KEAP1. On the other hand, Src kinase activity inhibition resulted in the release of p62-KEAP1 complex and led to KEAP1 association with NRF2, which can be therefore ubiquitinated and degraded (**Figure 4** and **Figure 5**). Our findings are in agreement with previous reports demonstrating that NRF2 and p62 can compete for KEAP1 binding (Komatsu *et al*, 2010; Jain *et al*, 2010; Lau *et al*, 2010). In addition, p62 is a major target of NRF2 transcription factor pointing to a positive feedback loop mechanism (Jain *et al*, 2010). Interestingly, we could demonstrate that the constitutive activation of Src relies on mTORC1 signaling to promote this signaling pathway (**Figure 6 and Supplementary S5-S6**). Our observations are consistent with recent studies that identified the aberrant activation of mTORC1 in GBM and in HCC cellular models as responsible for the upregulation of NRF2 (Komatsu *et al*, 2010; Pölönen *et al*, 2019). We can speculate that the aberrant activation of Src in cancer cells constitutively activates NRF2, in order to survive to unfavorable conditions. Importantly, we also demonstrated that Src activity promotes the expression of p62 (**Figure 3**) that when overexpressed, may further sustain mTORC1 signaling and NRF2 activation (Duran *et al*, 2011).

We reported that NRF2 acts as a downstream effector for Src deregulation in cancer, supporting cell proliferation and cell resistance to radiotherapy (**Figure 7**). Importantly, Src-dependent NRF2 activation may contribute to inhibit ferroptosis and indeed its targeting may increase sensitivity to ferroptosis, enhancing their efficacy in combination with IR (**Figure 8**).

The link between tyrosine kinases, NRF2 signaling and ferroptosis is still under-investigated. As NRF2 transcription factor controls several cellular responses, it will be interesting to perform transcriptomic analyses to further elucidate the panel of NRF2 target genes that may be modulated by Src kinase activity.

Our work suggests that the concomitant increased activity of Src and NRF2 may identify those tumors that may benefit from targeting this novel signaling axis. Of note, among GBM tumors, the mesenchymal subtype characterized by increased NRF2 expression (Pölönen *et al*, 2019) has also been shown to be more sensitive to Dasatinib treatment (Alhalabi *et al*, 2022). Since the constitutive activity of Src is a major feature of cancer, it will be interesting to extend the investigation to different tumors. Of note, being Src aberrantly induced downstream the activation of RTKs we speculate that RTKs may provide a general mechanism to impinge on NRF2 signaling independently of specific NRF2 or KEAP1 mutations. In summary, we provide the first evidence for a new connection between the aberrant activation of tyrosine kinases and the constitutive activation of NRF2 signaling and we identify an unexpected role for tyrosine phosphorylation signaling in the modulation of ferroptosis.

## METHODS

### Cell culture

U87MG and T98G (originally obtained by ATCC) were cultured in Dulbecco’s modified Eagle’s medium (DMEM) supplemented with 10% fetal bovine serum (FBS), 100 U/ml penicillin and 100 mg/ml streptomycin (Sigma-Alrich). GBMSC83 cells, a well-characterized mesenchymal GBM cellular model, were cultured as neurospheres in non-adherent conditions in DMEM/F12 supplemented by B27 Supplement (50x), EGF (20 ng/ml), and hβFGF (10 ng/ml) as previously described (Minata *et al*, 2019; Mao *et al*, 2013). All GBM cells were maintained in a humidified 5% CO, 37°C incubator and were routinely tested negative for mycoplasma contamination.

Stable cell lines, overexpressing Src *wild-type* or catalytically inactive mutant *kinase dead* in p-MX-psCESAR vector, were generated by retroviral infection followed by cell sorting for GFP^+^ cells. For transient transfection experiments cells were seeded the day before and transfected using polyethylenimine (PEI) (Tebu-Bio), following manifacturer’s instructions. Src constructs pSGT-Src-Y527F and pSGT-K295M and empty pSGT vector were previously described (Cursi *et al*, 2006).

### Antibodies and other reagents

Primary antibodies used are as follows: anti-NRF2 (D1Z9C) (Cell Signaling Technology), anti-p62 (SQSTM1) (PM045, MBL International), anti-phospho-p62 (SQSTM1) (Ser349) (PM074, MBL International), anti-Vinculin (E1E9V) (Cell Signaling Technology), anti-Lamin (E-1) (sc-376248, Santa Cruz Biotechnology), anti-GAPDH (D16H11) (Cell Signaling Technology); anti-Src (2108S) (Cell Signaling Technology), anti-phospho-Src (Tyr416) (2101S) (Cell Signaling Technology), anti-KEAP1 (G-2) (sc-365626, Santa Cruz Biotechnology), anti-Heme Oxygenase 1 (A-3) (sc-136960, Santa Cruz Biotechnology), anti-NQO1 (A-180) (sc-32793, Santa Cruz Biotechnology), anti-p70S6K (Cell Signaling Technology), anti-phospho-p70S6K (Thr389) (Cell Signaling Technology), anti-ERK 1/2 (137F5) (Cell Signaling Technology), anti-phospho-ERK 1/2 (Thr202/Tyr204) (D13.14.4E) (Cell Signaling Technology), anti-Ubiquitin (P4D1, Santa Cruz Biotechnology).

Dasatinib, Trigonelline, Rapamycin, Ferrostatin-1, Erastin and Cycloheximide were purchased from Sigma-Aldrich; PP2 (sc-202769) from Santa Cruz Biotechnology; ML385 from Selleck Chemicals.

### Fluorescence microscopy

Cells were seeded on coverslips and grown at 37 °C and 5% CO_2_. After treatments cells were washed with 1x phosphate buffer saline (PBS), fixed in 4% paraformaldehyde for 15 min at room temperature, permeabilized with PBS/Triton X-100 0.3% solution for 10 min, blocked with BSA 3% solution in PBS for 1 h and incubated overnight with primary antibodies (NRF2,1:50; KEAP1, 1:50; p62: 1:1000) in a humid chamber at 4 °C. Secondary antibodies (Thermo Fisher Scientific, 1:500) were applied for 1 h at room temperature, and nuclei were stained with Hoechst 33342 (Thermo Fisher Scientific) for 15 min. Images were acquired by Fluorescence microscopy (ZEISS) and processed with Fiji version 2.3.

Confocal microscopy experiments were performed by using LSM800 microscope (ZEISS) equipped with a 63x oil objective and with ZEN blue imaging software. Staining intensities were analyzed by using Fiji software (ImageJ) to obtain the nuclear/cytoplasmatic ratio. For colocalization studies, at least three slices were taken at a Z-stack distance of 0.3µm. Following acquisition, images were imported into Fiji for analysis and quantification of co-localizing particles between two channels by using the open-source plugin ComDet 0.5.2. Colocalization parameters: maximum distance between the center of two particles ≤ 4 pixels; particle size ≥ 3; intensity threshold = 5.

### Protein extracts, immunoprecipitation, nuclei/cytoplasm fractionation and immunoblotting analyses

Total cell extracts were prepared in Buffer A (10 mM Hepes [pH 7.9], 10 mM KCl, 1.5 mM MgCl_2_, 0.5 mM DTT, 0,1% NP-40, 5 mM EDTA, 5 mM EGTA, 1 mM phenylmethylsulfonyl fluoride, 25 mM NaF, 1 mM sodium orthovanadate,10mg/ml TPCK, 5mg/ml TLCK, 1mg/ml leupeptin, 10mg/ml soybean trypsin inhibitor, 1mg/ml aprotinin). Lysates were incubated for 20 min on ice, sonicated and centrifugated 12,000 g at 4 °C for 30 min.

For nuclei/cytoplasm cell fractionation, cells were lysate in the same Buffer A, without adding 0.1% NP-40. After incubation for 20 min on ice, NP-40 (0.1% final concentration) was added, and nuclei were harvested by centrifugation at 12,000 g at 4 °C for 30 s. The cytoplasmic fraction was recovered, and nuclear proteins were extracted from the pellet in Buffer A, completed with 0,05% NP-40 for 30 min on ice followed by sonication and centrifugation at 12,000 g at 4 °C for 30 s. For immunoprecipitation experiments total cell extracts were incubated with primary antibody for 3 hrs on a rotating wheel at 4 °C, followed by 45 min incubation with Protein G beads (Invitrogen). The complex was washed four times in ice cold PBS, denatured for 5 min at 95 °C. For immunoblotting, 20–50mg of proteins were separated by sodium dodecyl sulfate (SDS) polyacrylamide gel electrophoresis (PAGE), blotted onto nitrocellulose membrane, and detected with specific antibodies.

### Cell death analysis

Cell death was evaluated 48 hrs after ionizing radiation (10 Gy) by using a CytoFLEX S (Beckman Coulter, Milan, Italy) instrument. 1 × 10^6^ cells were collected, centrifuged at 4°C for 5 min at 300 × g and double-stained by using Annexin V-APC-propidium iodide (PI) kit, according to manufacturer’s instructions (eBioscienceTM Annexin V Apoptosis Detection Kits, ThermoFisher Scientific). For GFP^+^ cells, PI staining was replaced with DAPI. Unstained samples were used as control. Quality control was evaluated using CytoFLEX Daily QC Fluorospheres (Beckman Coulter).

FCS files were analyzed using CytExpert version 2.2 software (Beckman Coulter). Dead cells (Annexin V+/PI+ cells) were graphed as fold change to control conditions.

### Proliferation Assay

Cells were seeded in 12-well culture plate (1 x 10^5^ cells/ml) and incubated at 37 °C, 5% CO_2_. After 24 hrs cells were treated with Trigonelline 5μM. Cells were detached and counted 24, 48 and 72 hrs after treatment. Data were expressed as mean and SD. The assays were repeated four times.

### Clonogenic Survival Assay

Cells were plated in 6-well culture plate (1000 cells/well) and incubated at 37 °C, 5% CO2 for colony formation. After 10-15 days, colonies were fixed and stained with a solution with 10% (v/v) methanol and 0.5% Cristal Violet for 20 min for colony visualization. The stained colonies (>50 cells) were counted. Data were expressed as mean and SD. The assays were repeated four times.

### Real-Time PCR (RT–qPCR)

Cells were homogenized in TRI Reagent (Thermo Fisher Scientific), and RNA was extracted in accordance with manufacturer’s protocol. 1 mg of total RNA was retrotranscribed in cDNA using the SensiFAST cDNA Synthesis KIT (Bioline). Specific sets of primer pairs were designed and tested with primerBLAST (NCBI, see list below). RT–qPCR was performed using the SensiFAST Syber Low-ROX kit (Bioline) QuantStudio 3 RealTime qPCR (Applied Biosystems). Data were analyzed by using the second-derivative maximum method. The fold changes in mRNA levels were determined relative to a control after normalizing to the internal standard, Actin. Primers used are listed below:

### Gene Forward primer Reverse primer

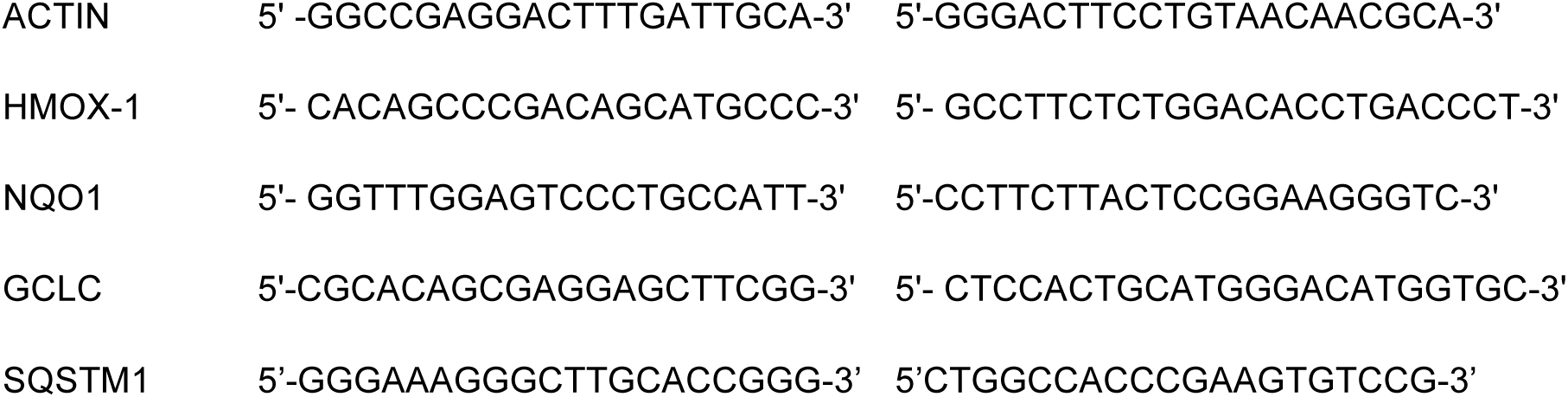
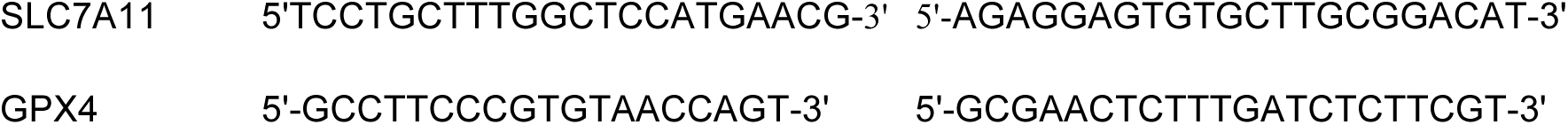

### Lipid peroxidation measurement

Membrane lipid peroxidation was evaluated 48 hrs after ionizing radiation (10 Gy), in the presence or not of Dasatinib and Ferrostatin-1. Cells were incubated for 30 min at 37°C with 5 mM of BODIPY665/676 probe (Thermo Fischer Scientific) dissolved in PBS, washed twice in PBS and analyzed by using a CytoFLEX S (Beckman Coulter, Milan, Italy) instrument. Unstained sample was used as negative control. FCS files were analyzed using CytExpert version 2.2 software (Beckman Coulter).

### Public available datasets

The expression profile of glioblastoma patients (GBM) was downloaded from TCGA Data Portal (https://tcga-data.nci.nih.gov) using the recommended GDC data transfer tool. The processed data (level 3) were used. To date, there are 163 patients with unrestricted for publication RNA-seq(V2) data. The expression data of normal brain tissues was downloaded from GTEx portal (V8) (https://gtexportal.org/home/) where a total of 3869 samples belonging to 13 different brain regions are available.

### Pathway variation analysis

Pathway enrichment score was calculated using Gene Set Variation Analysis tool (GSVA) (Hänzelmann *et al*, 2013). A matrix containing pathway enrichment scores for each gene set and sample was obtained using the TCGA and GTEx gene expression matrices and the MSigDB Halmark gene set collection (Subramanian *et al*, 2005). Differential pathway activity between GBM and normal brain cortex was calculated using Deseq2 (Love *et al*, 2014) and pathways with p-value<0.05 and |log2 FC| > 0.3 were selected as significant.

### Statistical analyses

All experiments were replicated at least three times (biological replicates) and data were presented as mean ± SD or ± SE, as indicated in the figure legends. The significance of the differences between populations of data were assessed according to the Student’s two-tailed t-test (independent samples) with level of significance of at least P ≤ 0.05.

## ACNOWLEDGMENTS

We thank Maria Pia Gentileschi for kindly providing technical support for irradiation experiments and all the members of our lab for critical reading of the manuscript and for helpful discussion.

This work has been supported by research grants from Associazione Italiana per la Ricerca sul Cancro AIRC-IG2016-n.19069, AIRC-IG2021-n.26230, and Italian Ministry of Health, RF-2016-02362022 to D.B.; C.Ci has been supported by a FIRC-AIRC fellowship for Italy “Filomena Todini”; C.Co. work also been supported by AIRC IG2016-n.19069 and AIRC-IG2021-n.26230, I.T. is supported by a MUR fellowship to the PhD Program in Cellular and Molecular Biology, Department of Biology, University of Tor Vergata.

## AUTHOR CONTRIBUTIONS

C.Ci., designed and performed most of the experiments, contributed to data analysis, interpretation and to write the article; I.T. contributed to perform experiments; C.Co contributed to perform experiments on protein stability; M.D. and G.B. perform cytofluorimetry analyses to sort cell lines stably overexpressing Src constructs; G.P. and M.H.C. performed bioinformatic analyses; D.B. designed the experiments, evaluated the data and wrote the paper.

## Supplementary Figures

**S1.**
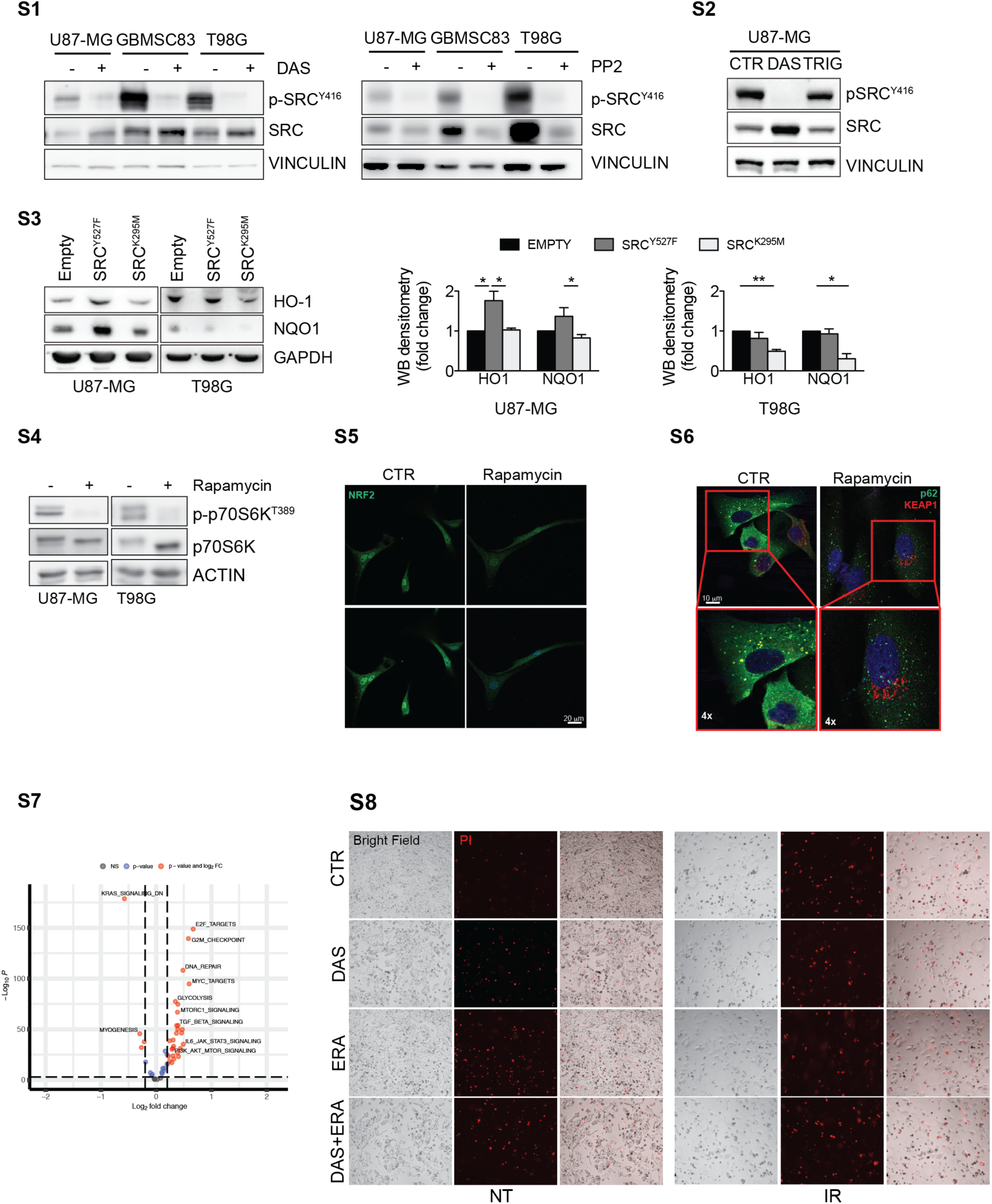
Immunoblotting of p-Src^Y416^ and Src in U87-MG, T98G and GBMSC83 cells treated with DAS 10nM (U87-MG and T98G) and 1μM (GBMSC83) (left) or PP2 5μM (right) for 24hrs. Vinculin was used as loading control. **S2**. Immunoblotting of p-Src^Y416^ and Src in U87-MG cells treated with DAS 10nM or Trigonelline 5μM for 16hrs. **S3**. Immunoblotting and relative densitometric analysis of NRF2 targets, HO-1 and NQO1, in U87-MG and T98G cells after 24hrs of transient transfections with active Src (Src^Y527F^) or catalytically inactive mutant (Src^K295M^). **S4**. Immunoblotting of p-p70S6K^T389^ and p70S6K in U87-MG and T98G cells treated with Rapamycin 100nM for 48hrs. Actin was used as loading control. **S5**. Immunofluorescence staining in U87-MG cells treated with Rapamycin 100nM for 48hrs. NRF2 (green); DNA (Hoechst, blu). **S6**. Confocal microscopy analyses of U87-MG cells treated with Rapamycin 100nM for 48hrs. p62 (green); KEAP1 (red); DNA (Hoechst, blue); 4x digital magnification showing merged signals. **S7**. Volcano plot. The log2 FC indicates the mean activity level for each pathway. Each dot represents one pathway. Black and blue dots represent pathways with no significant p-value and no significant FC respectively, red dots represent up-and down-regulated pathways in GBM (TCGA data) compared with normal brain cortex (GTEx data); **S10**. Rapresentative Bright Field images and propidium iodide (PI^+^, red) staining in T98G cells after 48hrs irradiation (10Gy), with or without DAS 10nM and/or Erastin (ERA) 1μM. Statistical analyses: paired (**S3**) and unpaired (**S9,S10**) Student’s t-test: (* p < 0.05; ** p < 0.01).

